# Two Genomes, One Metabolome: Mitonuclear Incompatibility Remodels Developmental Metabolism and Fitness in *Drosophila*

**DOI:** 10.64898/2026.05.29.728848

**Authors:** Akansha Singh, Brian Frias, Briana M. Deaver, Ariana C. Edwards, Jordy K. Cambara, Omera B. Matoo

**Author notes:** Corresponding author: Omera B. Matoo, 414 E Clark St., 170A, Churchill-Haines Lab, Vermillion, SD - 57069.

## Abstract

Mitochondrial-nuclear (hereafter mitonuclear) genetic variation can alter cellular bioenergetics and metabolism via jointly encoding the subunits of oxidative phosphorylation (OXPHOS) system. Here we tested whether genetic variation in biochemical and bioenergetic phenotype can scale up across higher levels of biological organization to affect organismal development. We used a panel of *(mtDNA); nDNA* genotypes created by asymmetric substitution of divergent mtDNA between *Drosophila melanogaster* and its sister species *D. simulans*. Two genotypes – *(ore); OreR* and *(ore); Aut* – carry coevolved mitonuclear genomes from *D. melanogaster*, whereas the other two – *(w^501^); Aut* and *(w^501^); OreR* – harbor *D. simulans* mitochondrial DNA introgressed onto *D. melanogaster* nuclear backgrounds. We utilized untargeted metabolomics to track down the comprehensive biochemical footprint of the joint genomic architecture in the metabolome of these genotypes. We show that in *(w^501^); OreR* larvae with mitonuclear genome incompatibility there is extensive and coordinated metabolic remodeling of carbon, nitrogen, and redox balance, characterized by increased glycolysis, limited TCA cycle, amino acid scarcity, reduced nucleic acid balance, lipid remodeling, with compensatory activation of antioxidant pathways and possible epigenetic modulation by metabolites. This biochemical rewiring puts a physiological constraint that prioritizes maintenance metabolism over larval growth, leading to reduced body size, slow locomotion and delayed development, despite a compensatory increase in feeding in these larvae. These findings underscore the context dependency of mitonuclear interactions, which can uniquely scale up to influence organismal energy budget trade-offs, fitness and life-history evolution.

## Introduction

Metabolism is an emergent property of biochemical reactions within an organism that mediate the continuous acquisition, transformation, and utilization of energy (Glazier 2015). This energy is subsequently allocated into maintenance, growth, activity and reproduction (Sokolova 2013). Trade-offs in energy allocation among these functions can have important fitness implications (Sokolova 2013), particularly during development of holometabolous insects, including *Drosophila*, which are characterized by rapid and massive larval growth (Church and Robertson 1966; Miyazawa and Aulehla 2018). Because development is energetically demanding, mitochondria - the cellular energy harvesters - are integrated tightly into development rather than being evolutionary bystanders (Jacobs et al. 2020; Liu and Auwerx 2023).

Mitochondrial energy transduction is mediated by the electron transfer system (ETS), which generates ATP via oxidative phosphorylation (OXPHOS), reactive oxygen species (ROS) for signaling, and dissipates heat (Gray et al. 1999). Additionally, three cellular pathways (glycolysis, oxidation of fatty acids and breakdown of amino acids) converge on the tricarboxylic acid (TCA) cycle in the mitochondria (Anderson et al. 2019; Jacobs et al. 2020). In rapidly growing *Drosophila* larvae these reactions, together with NADPH as the reducing power, enable conversion of carbon backbones derived from food into new biomass for growth (Tennessen and Thummel 2011; Miyazawa and Aulehla 2018; Jacobs et al. 2020). Impaired ETS constrains flux through the TCA cycle, affecting bioenergetics, signaling, growth, and development (Tennessen and Thummel 2011; Anderson et al. 2019; Martínez-Reyes and Chandel 2020; Liu and Birsoy 2023). For example, studies have demonstrated that mitochondria regulates the segmentation clock in mice and humans and acts as a pacemaker of species-specific development through its control of cellular energetics and redox homeostasis (Diaz-Cuadros et al. 2023; Iwata et al. 2023).

Substantial genetic variation, even within species, exists in metabolic enzymes due to differences at enzyme-coding loci, *trans*-acting, and epistatic variation throughout the genome (Hubby and Lewontin 1966; Clark et al. 1995; Montooth et al. 2003). OXPHOS complexes are mosaics of polypeptides jointly encoded by two cellular genomes: nuclear DNA (nDNA) and mitochondrial DNA (mtDNA) (Gray et al. 1999). The mtDNA encodes 13 OXPHOS proteins, which are synthesized on the organelle’s own translational machinery, whereas the rest of mitochondrial proteins (∼1200) are encoded by nDNA, synthesized in the cytoplasm, and then imported into the mitochondria (Kummer and Ban 2021). Variation in metabolic phenotype is, therefore, associated with variation in both nuclear, and mitochondrial genomes, and with epistatic interactions between them (GXG) (Hill 2015 and references therein). Given the importance of metabolism to support maintenance, growth, activity, and reproduction, it is not surprising that both positive and purifying selection shape variation and divergence in mitochondrial and nuclear (henceforth mitonuclear) genomes (Rand and Mossman 2020; Camus and Dhawanjewar 2023). Discordance between mitonuclear genomes can impair OXPHOS, resulting in metabolic inefficiency and fitness defects, including in development (Meiklejohn et al. 2013; Buchanan et al. 2018; Matoo et al. 2019; Pichaud et al. 2019; Ghiselli and Milani 2020; Burton 2022; Camus et al. 2023). Despite this, our understanding of integrated molecular mechanisms that maintain biochemical-physiological-evolutionary communication and (co)adaptation between these two genomes remains incomplete.

Previous studies have shown that *Drosophila* L2 (second instar) larvae with mitonuclear genome incompatibility are bioenergetically compromised, exhibit slower growth and have a two-day delay in development compared to their matched genetic controls (Meiklejohn et al. 2013; Matoo et al. 2019). In the present study, we test whether mitonuclear interactions (1) mediate any remodeling of the internal metabolic state, and (2) scale up the hierarchical levels of biological organization to affect larval traits (specifically body size, locomotion and feeding) that determine fitness. We hypothesize that the larvae with mitonuclear incompatibility will exhibit extensive metabolic remodeling and reduced whole-organism fitness, reflecting a shift in allocation toward survival rather than growth and development. These findings are broadly relevant to understand whether genetic variation underlying metabolic phenotypes may be adaptive, deleterious, nearly neutral in isolation but can exhibit strong fitness effects due to mitonuclear epistasis and different environmental contexts (GXGXE).

## Methods

### Fly Stocks and Mitonuclear Panel

We used the *Drosophila* mitonuclear panel previously generated by Montooth et al. 2010 (Figure S1). Divergent mtDNAs from closely related sister species *Drosophila melanogaster* and *Drosophila simulans* were asymmetrically substituted to create four novel (*mtDNA); nDNA* genotypes: [1] *(ore); OreR*, [2] *(ore); Aut*, [3] *(w^501^); Aut* and [4] *(w^501^); OreR*. Individuals from *(ore); OreR* and *(ore); Aut* have coevolved mitonuclear genomes from *D. melanogaster* whereas individuals from *(w^501^); Aut* and *(w^501^); OreR* have mismatched genomes, with mtDNA introgressed from *D. simulans*. In the mismatched *(w^501^); OreR* genotype, individuals have a genetic incompatibility between naturally occurring single nucleotide polymorphisms (SNPs) in the mt-tRNA^Tyr^ gene and the nuclear-encoded mt-tyrosyl-tRNA synthetase gene *Aatm* that aminoacylates the mt-tRNA^Tyr^, putatively compromising their protein translation (Montooth et al. 2010). The remaining three genotypes *(ore); OreR*, *(ore); Aut* and (*w^501^); Aut* serve as matched genetic controls.

### Sampling

Flies from all genotypes were raised in incubators (Shel Lab, USA) on Nutri-Fly^®^ Molasses formulation (Genesee Scientific, USA), acclimated to 25 °C, 65% relative humidity under a 12:12 h light: dark cycle and allowed to lay eggs overnight on fresh food supplemented with active yeast paste. L2 larvae were collected based on developmental time and distinguishing morphological features (Ashburner 1989) under Leica S9i stereomicroscope (Leica Microsystems, Germany) equipped with an in-built 10 MP CMOS digital camera (6.1X-55X magnification) rinsed in deionized water and then 1X phosphate buffered saline (PBS) to remove any food particles.

### LC-MS Based Untargeted Metabolomics

For each genotype, five biological replicates were collected, with five L2 larvae pooled per replicate. Samples were collected as described above, flash-frozen and stored at -80 °C and subsequently shipped to The Center for Mass Spectrometry and Metabolic Tracing Core Facility (Washington State University, St. Louis) for LC-MS based whole-body tissue untargeted metabolomics. Briefly, larvae were homogenized using an Omni Bead Ruptor Elite homogenizer (Omni International, USA). Samples were extracted with methanol: acetonitrile: water (2:2:1, v/v/v) at a ratio of 40 µL of extraction solvent per 1 mg of tissue. Homogenization was performed for two cycles at 6 m/s for 30 s per cycle. Extracts were incubated at -20 °C for 1 h to facilitate protein precipitation. Following incubation, samples were centrifuged at 14,000 × g for 10 min at 4 °C. The supernatant was transferred to LC-MS vials, and polar extracts were stored at -80 °C until analysis. Ultra-high performance liquid chromatography coupled with mass spectrometry (UHPLC/MS) analyses were conducted using a Thermo Scientific Vanquish Flex UHPLC system, interfaced with a Thermo Scientific Orbitrap ID-X Mass Spectrometer. For the separation of polar metabolites, a HILICON iHILIC-(P) Classic HILIC column (100 x 2.1 mm, 5 µm) with a HILICON iHILIC-(P) Classic guard column (20 x 2.1 mm, 5 µm) was utilized. The mobile-phase solvents consisted of solvent A = 20 mM ammonium bicarbonate, 2.5 µM medronic acid, 0.1% ammonium hydroxide in 95:5 water: acetonitrile and solvent B = 95:5 acetonitrile: water. The column compartment temperature was maintained at 45 °C, and metabolites were eluted using a linear gradient at a flow rate of 250 µL/min as follows: 0-1 min, 90% B; 12 min, 35% B; 12.5-14.5 min, 25% B; 15 min, back to 90% B. Data was acquired in both positive and negative ion mode. The LC-MS data were processed and analyzed using XCMS, Compound Discoverer, and Skyline (Adams et al. 2020).

### Organismal Traits

Body size measurements were performed using protocol from Garcia et al. (2025). L2 larvae (n = 25/genotype) were transferred to a glass dish mounted on the stage of a stereomicroscope. Individual larvae were immersed in deionized water at 80 °C for ∼5 s to achieve immobilization and body distension. Afterwards, larvae were blotted dry and imaged. Images were analyzed using FiJi/ ImageJ v1.54p (NIH, USA) (Schneider et al. 2012), where larval size was estimated by measuring body length and width (mm) along the longitudinal and transverse axes. Crawling assay was adapted from Garcia et al. (2025). L2 larvae (n = 50 /genotype) were placed on 2% agar with a mm grid scale (0.2 cm²). Larvae were acclimated for 5 min, following which the speed of crawling/locomotion was quantified as the number of lines crossed by individual larva per minute. Peristaltic propagation (laterally symmetric waves of muscular contraction and relaxation from anterior to posterior end of the larva) that induces crawling was recorded per minute for each larva. Food consumption assays were adapted from Kaun et al. (2007) and Garcia et al. (2025). L2 larvae (n = 50 larvae/genotype) were placed in petri dishes lined with tissue paper saturated with 1X PBS for a 3 h fasting period. Larvae were then transferred to 2% agar plates coated with freshly prepared yeast paste (32.5% w/v in water) containing Brilliant Blue R-250 (0.7% w/w) and allowed to feed for 10 min at 25 °C. Following feeding, larvae were immobilized, imaged, and food intake was quantified in FiJi/ImageJ v1.54p (NIH, USA) (Schneider et al. 2012) by measuring the blue-stained area relative to total larval body area.

### Statistical Analysis

192 metabolites across the four genotypes were detected, generating a peak-intensity data matrix that was imported into MetaboAnalyst 6.0 (Pang et al. 2024). Data were analyzed using a hierarchical framework integrating exploratory, predictive, univariate, multivariate, and systems-level approaches (Ren et al. 2015). Dimensionality reduction was performed using Principal Component Analysis (PCA) to visualize global metabolic patterns, while Pearson correlation-based hierarchical clustering was used to further explore relationships among samples. To assess whether metabolomic divergence was predictive and robust, a supervised machine-learning approach, Random Forest (RF), was implemented using the 192 detected metabolites as predictors. RF is an ensemble approach that combines multiple uncorrelated decision trees generated via bootstrap aggregation (bagging), improving robustness to noise and outliers. Univariate analyses included one-way Analysis of Variance (ANOVA; genotype as factor), two-way ANOVA (nDNA and mtDNA as factors), fold-changes, and targeted pairwise comparisons. To account for the multivariate structure of metabolomics, partial least squares–discriminant analysis (PLS-DA), and variable importance in projection (VIP) scores were used to identify metabolites contributing to differences among genotype-pairs. Previous studies have shown that, for complex datasets, PCA alone can lead to overfitting, and incorporating prior experimental design information improves estimation of the underlying structure (Bertinetto et al. 2020). To address this, we used a multivariate ANOVA-simultaneous component analysis (ASCA), which decomposes the metabolite data into effect matrices corresponding to experimental factors (nDNA, mtDNA, and nDNA × mtDNA interaction) and a residual matrix (Bertinetto et al. 2020) using R v 4.3.2 (R Core Team 2025). PCA was then applied to each effect matrix to extract variation associated with each factor. Finally, to link metabolite changes to biological processes, we performed metabolite set enrichment analysis (MSEA) and pathway analysis using the Kyoto Encyclopedia of Genes and Genomes (KEGG) database with *Drosophila melanogaster* as the reference (KEGG:dme) in MetaboAnalyst 6.0. Pathway analysis integrates over-representation analysis (ORA) using Fisher’s Exact Test with topology-based network analysis based on relative betweenness centrality to evaluate both the statistical enrichment of metabolites within pathways (False Discovery Rate (FDR) ≤ 0.05) and their relative positions within metabolic networks (impact score). Prior to all statistical testing, metabolite peak intensities were normalized by sample sum, log2 (or log10 for pairwise comparisons) and autoscaled for all analyses (Figure S2).

Differences among genotypes in body size, crawling and feeding were evaluated using generalized linear models (GLMs) (Neuhaus and McCulloch 2011) with gamma distribution and log link in R v4.3.2 (R Core Team 2025). We derived multiplicative effect sizes from model coefficients by exponentiating parameter estimates (exp (coef)), which provide estimates of proportional change (i.e., fold differences) relative to the reference genotype (designated as *(ore); OreR*). To visualize variation in all traits, we calculated estimated marginal means (EMMs; Searle et al. 1980) with confidence intervals. A hierarchical model set for each organismal trait was also constructed to test the effects of nDNA, mtDNA, and their additive and interactive combinations, with model support assessed using AIC, ΔAIC, and likelihood ratio tests. For body size, we additionally used bootstrapping to estimate confidence intervals and assess robustness of parameter estimates.

## Results

### Exploratory Analysis of Metabolome

The two principal components in PCA cumulatively accounted for ∼53.70% (PC1 43.10%, PC2 ∼10.60%) of total variance (Figure 1A). Clear separation of *(w^501^); OreR* along PC1 was observed whereas variation among other three genotypes was comparatively smaller and partially overlapping. Hierarchical clustering of samples showed that *(w^501^); OreR* formed a distinct cluster with high within-group correlations, whereas the other three genotypes were more interspersed (Figure 1B). For the RF model, after n_tree_ = 500, the cumulative OOB error rates decreased to 0.20 (Figure 1C). Genotype-specific OOB error was 0.20 for *(ore); OreR*, 0.20 for *(ore); Aut*, 0.40 for *(w^501^); Aut*, and 0.00 for *(w^501^); OreR*, suggesting partial overlap in metabolic profiles among the first three genotypes, while *(w^501^); OreR* forms a separate group that is consistently classified without error.

**Figure 1.**
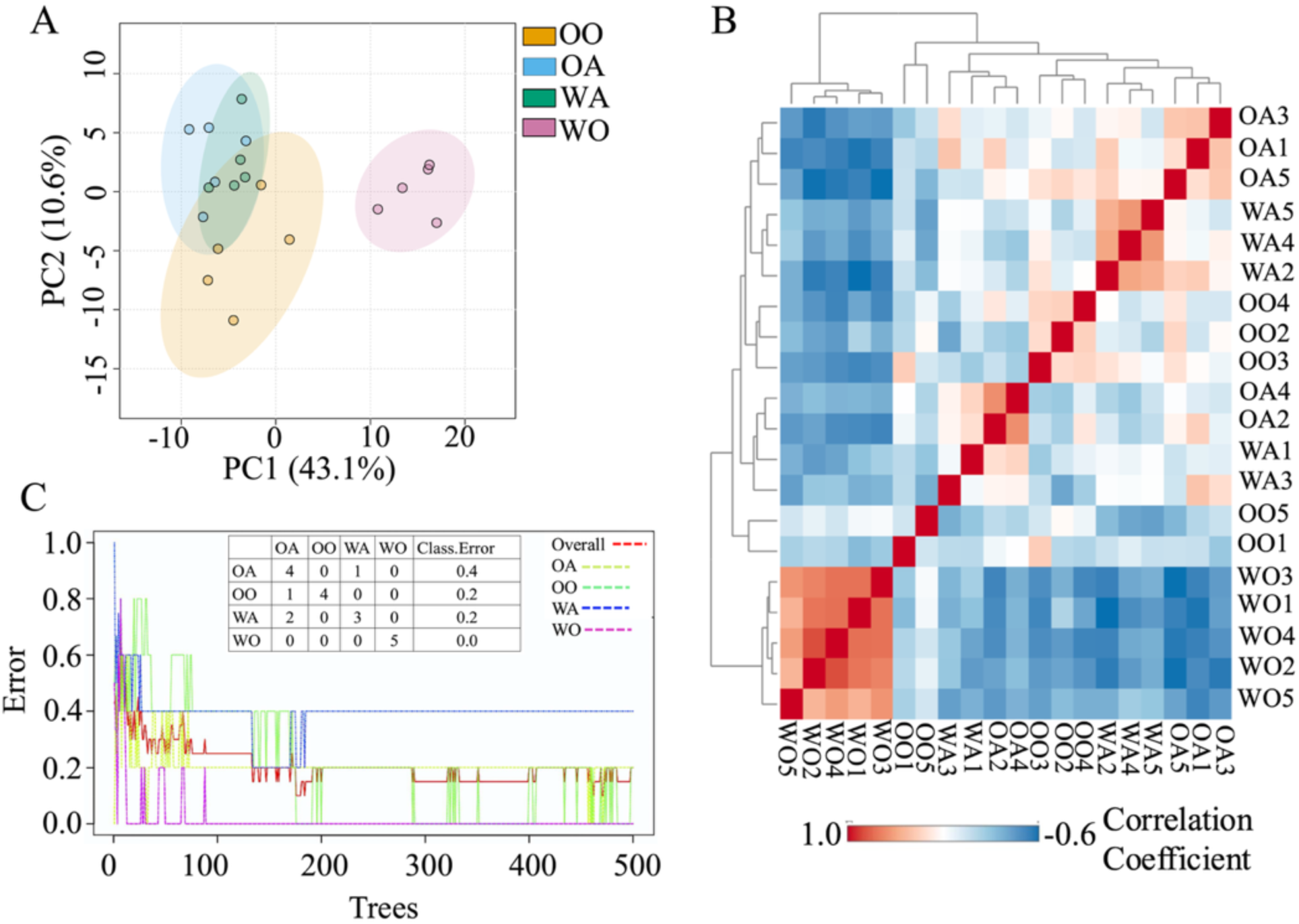
Exploratory Analysis and Predictive Modeling of Metabolome. [A] Principal Component Analysis score plot, [B] Pearson correlation-based hierarchical clustering across all samples, [C] Predictive classification of genotypes with a supervised Random Forest showing (right) stabilization graph for n=500 decision trees used to build the model and (inset) out-of-bag (OOB) error table. OO = *(ore); OreR*, OA = *(ore); Aut*, WA = (*w^501^); Aut*, WO = *(w^501^); OreR*. N = 5 biological replicates per genotype.

### Genotype Effect on the Metabolome

Genotype had a significant effect on metabolite abundance. One-way ANOVA identified 133 metabolites (69.27% of 192 detected) that differed significantly among genotypes (FDR *P*-adj ≤ 0.05; SI T1). From these, we specifically identified the subset of metabolites that were significantly less abundant (n = 45, 33.83%) and significantly more abundant (n = 45, 33.83%) in *(w^501^); OreR* only relative to all other three genotypes (SI T2). These differential changes in central carbon, amino acids, nucleic acids, lipids, and redox metabolism for 63 metabolites were visualized using the iPath3.0 (https://pathways.embl.de/ipath3.cgi) (Figure 2).

**Figure 2.**
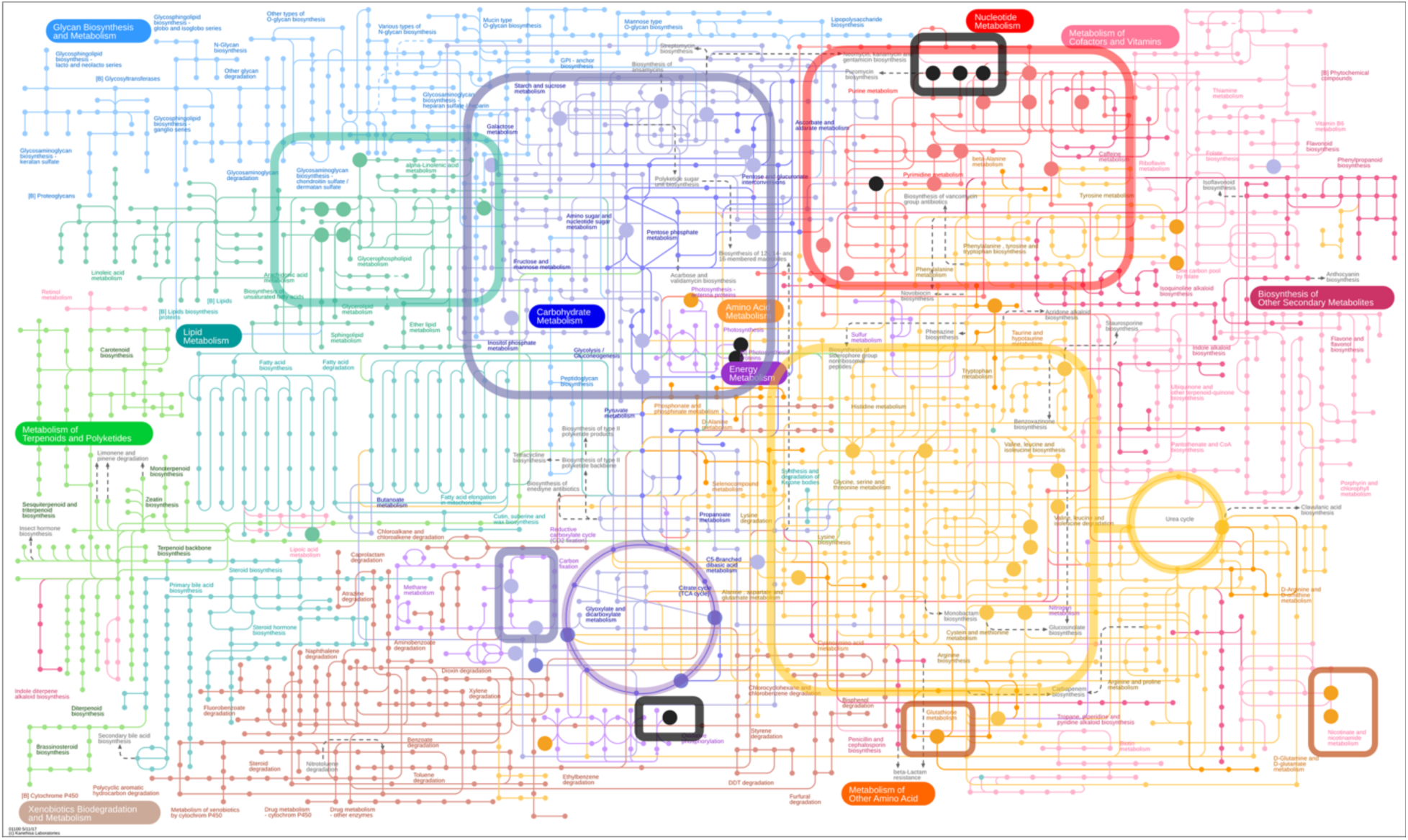
Differentially Abundant Metabolites in larvae with Mitonuclear Incompatible *(w^501^); OreR* Genotype in iPath3.0. iPath3.0 provides a comprehensive view of metabolic shifts by integrating upstream and downstream metabolite interactions and enzymatic reactions. The figure illustrates representative depleted amino acids (yellow), nucleic acids (red), TCA cycle intermediates (circled dark purple) and elevated lipid (green), redox (orange), currency (black) and phosphorylated sugar (boxed light purple) metabolites. Currency metabolites = ATP/ADP/AMP, NAD(H), NADP(H), FAD(H_2_).

We further performed pair-wise comparisons to identify (i) metabolites that change significantly between genotypes, (ii) metabolites that contribute most to overall genotype pair separation, and (iii) significant biological pathways that differ between genotype pairs.

#### a. Metabolome comparison among genotypes with co-evolved genomes

Among genotypes with shared co-evolved mtDNA (*ore*), different nDNA backgrounds (*OreR* vs. *Aut*) showed limited metabolic divergence. Only seven metabolites (|log_2_FC| ≥ 1, *P*-adjusted ≤ 0.05) were significantly more abundant in *(ore); Aut* compared to *(ore); OreR* (Figure 3A, SI T3). PLS-DA also identified these seven metabolites as having high discriminatory power (VIP ≥ 1.5), in addition to other contributing metabolites (Figure S3A). MSEA showed no significant pathways after FDR correction. However, several pathways, primarily associated with amino acid metabolism, were significant based on unadjusted *P*-values, including phenylalanine, tyrosine and tryptophan biosynthesis (*P* = 0.005, FDR = 0.42), phenylalanine metabolism (*P* = 0.01, FDR = 0.42), ubiquinone and other terpenoid-quinone biosynthesis (*P* = 0.02, FDR = 0.70), valine leucine and isoleucine degradation (*P* = 0.05, FDR = 0.88) and tyrosine metabolism (*P* = 0.05, FDR = 0.88).

**Figure 3.**
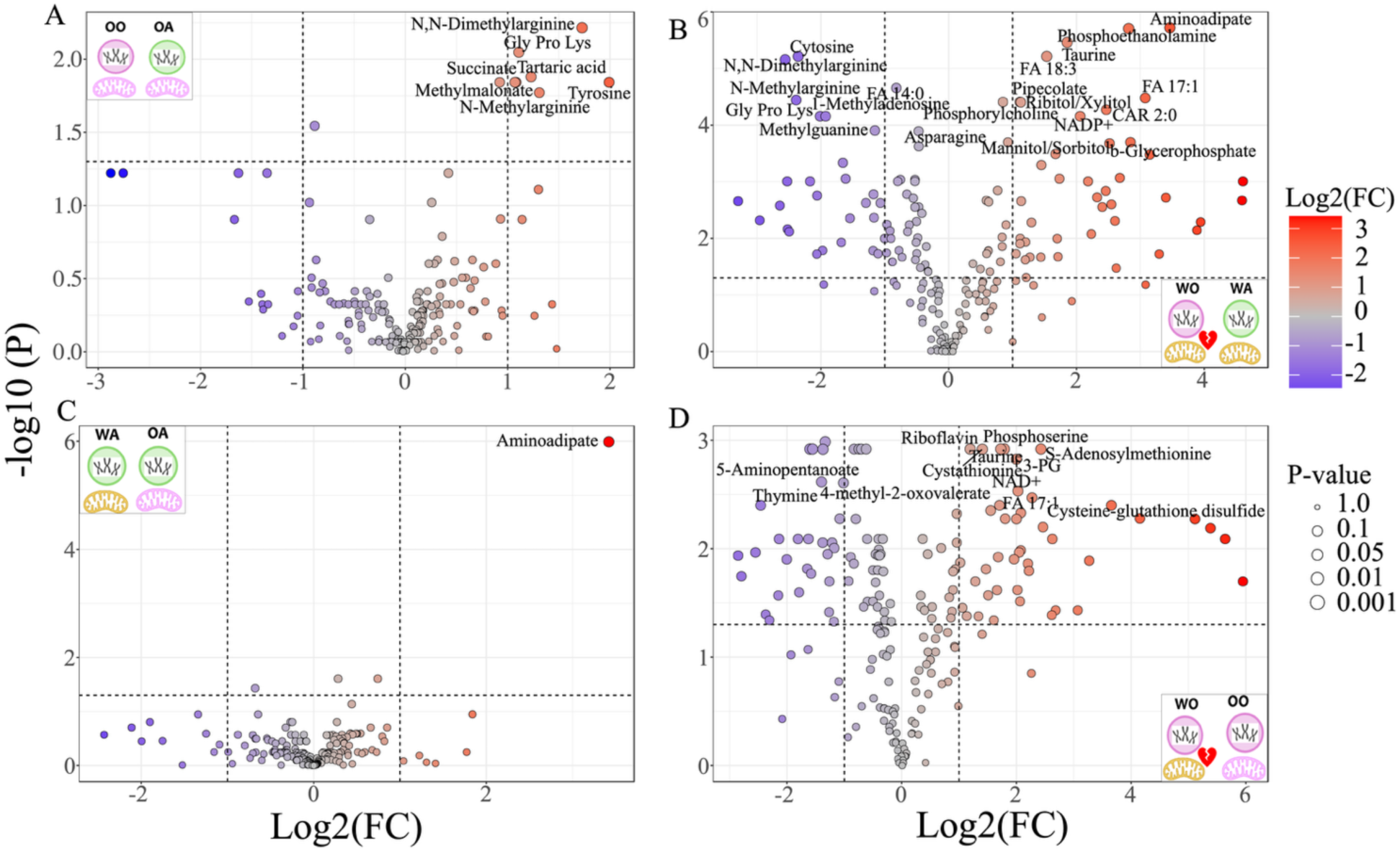
Differential Abundance Analysis of Metabolites in Genotype Pairs. Volcano plots depicting top 20 significantly up (red) and down (purple) metabolite pools [A] among coevolved genotypes OO vs OA, [B] among mismatched genotypes WO vs WA, [C-D] between coevolved vs. mismatched genotypes (WA vs. OA and WO vs. OO). OO = *(ore); OreR*, OA = *(ore); Aut*, WA = *(w^501^); Aut*, WO = *(w^501^); OreR*.

#### b. Metabolome comparison among genotypes with mismatched genomes

Among genotypes with shared but mismatched mtDNA (*w^501^*), different nDNA backgrounds (*OreR* vs. *Aut*) showed strong metabolic divergence. A total of 68 metabolites were significantly altered, with 41 metabolites significantly more abundant and 27 significantly less abundant in *(w^501^); OreR* relative to *(w^501^); Aut* (|log₂FC| ≥ 1, *P*-adjusted ≤ 0.05) (Figure 3B, SI T3). PLS-DA also identified 21 of these 68 metabolites with high discriminatory power (VIP ≥ 1.5), in addition to other contributing metabolites (Figure S3B). MSEA identified glycerophospholipid metabolism (*P* = 2.04 × 10^-^⁵, FDR = 0.0016), purine metabolism (*P* = 2.7 × 10^-^⁴, FDR = 0.0077), pyrimidine metabolism (*P* = 3.4 × 10^-^⁴, FDR = 0.0077) and 1C-pool by folate (*P* = 3.8 × 10^-^⁴, FDR = 0.0077) as significant pathways.

#### c. Metabolome comparison between co-evolved vs. mismatched genotype

While *(w^501^); Aut* had a conserved metabolic profile similar to *(ore); Aut*, the mitonuclear incompatible *(w^501^); OreR* genotype exhibited a significant metabolic divergence relative to *(ore); OreR*. For *(w^501^); Aut* vs. *(ore); Aut* comparison, aminoadipate was the only significantly different metabolite (|log₂FC| ≥ 1, *P*-adj < 0.05), and it was less abundant in *(w^501^); Aut*. MSEA associated aminoadipate with lysine degradation; however, this pathway was not significant after FDR correction (FDR = 1) (Figure 3C, SI T3). PLS-DA identified aminoadipate as having high discriminatory power (VIP ≥ 1.5), along with other contributing metabolites (Figure S3C). In contrast, *(w^501^); OreR* vs. *(ore); OreR* comparison revealed a total of 70 significantly altered metabolites, with 42 metabolites more abundant and 28 less abundant in *(w^501^); OreR* relative to *(ore); OreR* (|log₂FC| ≥ 1, *P*-adjusted ≤ 0.05) (Figure 3D, SI T3). PLS-DA with a VIP ≥ 1.0 identified a large number of discriminatory metabolites (n = 108); however, none met the more stringent criterion of VIP ≥ 1.5 applied in previous comparisons (SI T3). Instead, a subset of 17 metabolites exhibited intermediate VIP scores (≈1.30 -1.38), of which 12 overlapped with the 70 differentially abundant metabolites (Figure S3D). MSEA identified 1C-pool by folate (*P* = 0.0001, FDR = 0.0068), cysteine and methionine metabolism (*P* = 4.78E^-5^, FDR = 0.0038), and pyrimidine metabolism (*P* = 0.0012, FDR = 0.032) as significant pathways.

### nDNA and mtDNA Effects on the Metabolome

nDNA, mtDNA as well as their interaction (nDNA × mtDNA) significantly shaped the metabolome. Of the 192 metabolites, two-way ANOVA identified 139 (72.39%) as significantly affected by genomic factors (FDR *P*-adj ≤ 0.05) (Figure 4A). Among the 139 metabolites, 19 were associated with nDNA, 27 with mtDNA, 34 with both nDNA and mtDNA independently (additive effect) and 59 with nDNA × mtDNA interactions (mitonuclear epistasis) (Figure 4A, SI T4). A heatmap of the top 50 differentially abundant metabolites highlights these effects (Figure 4B). Metabolites identified as significant for nDNA, mtDNA, and their interaction (nDNA × mtDNA) were prioritized for pathway analysis. Based on a pathway impact threshold score > 0.09, nine pathways were initially identified for nDNA, seven for mtDNA, and 16 for the interaction. Although none remained significant after FDR correction, six (nDNA), three (mtDNA), and eight (interaction) pathways had unadjusted *P* ≤ 0.05 within this high-impact group.

**Figure 4.**
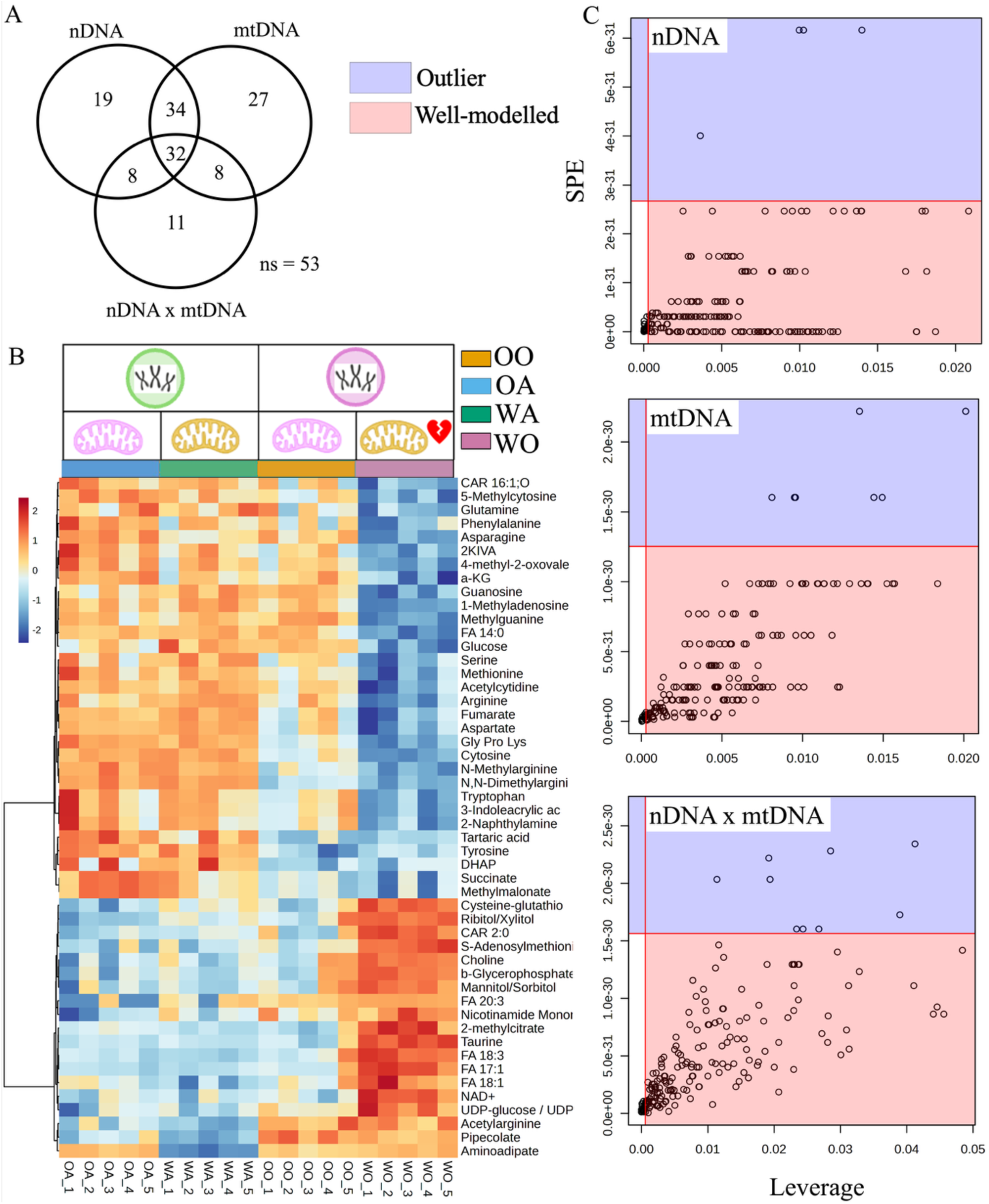
Effects of Nuclear and Mitochondrial Genomes on the Metabolome. (A) Venn diagram illustrating significant metabolites affected by nDNA, mtDNA, and their interaction. (B) Top 50 differentially abundant metabolites identified by two-way ANOVA across all four genotypes. (C) ANOVA-Simultaneous Component Analysis (ASCA) SPE (squared prediction error) –leverage plots corresponding to the three effect matrices: nDNA, mtDNA, and nDNA × mtDNA interaction. Each plot identifies well-modeled metabolites (pink) and outliers (purple) within each ASCA component.

We further performed ASCA to partition total variation into factor-specific effects (nDNA, mtDNA) and their interaction (nDNA × mtDNA). These models explained 22.68%, 18.66%, and 12.43% of the total variance, respectively. Permutation testing (1,000 permutations) confirmed that all effects were significant (nDNA: *P* = 0.000; mtDNA: *P* = 0.002; nDNA × mtDNA: *P* = 0.034) (Figure S4A). Score plots revealed clear separation of nDNA (*OreR* vs. *Aut*) and mtDNA (*ore* vs. *w^501^*) backgrounds along the first component, which explained ∼100% of the variation for each main effect (Figure S4B). In contrast, the interaction score plot exhibited a crossover pattern, with the direction of mtDNA effects reversing across nDNA backgrounds (Figure S4B), indicating context dependency of mitonuclear epistasis. Model performance was evaluated using leverage (threshold = 0.2) and squared prediction error (SPE, *a* = 0.05). Specifically, the nDNA, mtDNA, and interaction (nDNA × mtDNA) models adequately captured 166, 176, and 165 metabolites, respectively (Figure 4C). Metabolites exhibiting high leverage (> 0.01) and low SPE, indicative of strong influence and good model fit, showed substantial overlap with those identified as significant in the 2-way ANOVA.

### Genetic Architecture of Organismal Traits

#### Body Size

Genotype significantly affected *OreR* larvae consistently smaller than the other three genotypes. These differences were largely explained by additive effects of nDNA and mtDNA. GLM-based multiplicative effect sizes indicated that, relative to the reference *(ore); OreR*, *(ore); Aut* larvae were ∼9.2% longer (exp(β) = 1.09, *P* = 0.0001), *(w^501^); Aut* did not differ (exp(β) = 0.99, *P* = 0.70), and *(w^501^); OreR* were ∼7.5% shorter (exp(β) = 0.92, *P* = 0.0008) (Figure 5A-B, SI T5). Tukey-adjusted pairwise comparisons of EMM further showed that *(w^501^); OreR* larvae were significantly shorter than both *(ore); Aut* (*P* < 0.0001) and *(w^501^); Aut* (*P* = 0.015). Model comparisons (AIC, ΔAIC, and likelihood ratio tests) supported additive model of nDNA and mtDNA (*P* = 2.33E^-07^) (Table 1), with *Aut* nDNA increasing length relative to *OreR* and *ore* mtDNA increasing length relative to *w^501^* (both *P* < 0.05) (Figure 5B). The nDNA × mtDNA interaction was not significant (*P* = 0.64) and showed low bootstrap stability (6.5% of iterations significant). A similar pattern was observed for width. Relative to *(ore); OreR*, *(ore); Aut* larvae were 13% wider (exp(β) = 1.12, *P* = 1.93E^-06^), whereas *(w^501^); Aut* were 5.3% narrower (exp(β) = 0.95, *P* = 0.02) and *(w^501^); OreR* were 13.2% narrower (exp(β) = 0.87, *P* = 8.32E^-08^) (Figure 5C-D, SI T5). Pairwise comparisons confirmed that *(w^501^); OreR* larvae were significantly less wide than both *(ore); Aut* (*P* < 0.0001) and *(w^501^); Aut* (*P* = 0.0031). Model comparisons supported additive effects of nDNA and mtDNA on width (*P* = 4.70E^-06^) (Table 1), with *Aut* nDNA and *ore* mtDNA both increasing width (*P* < 0.001) (Figure 5D), while the interaction term remained non-significant (*P* = 0.40) and unstable under bootstrap resampling (1.9% significant iterations).

**Figure 5.**
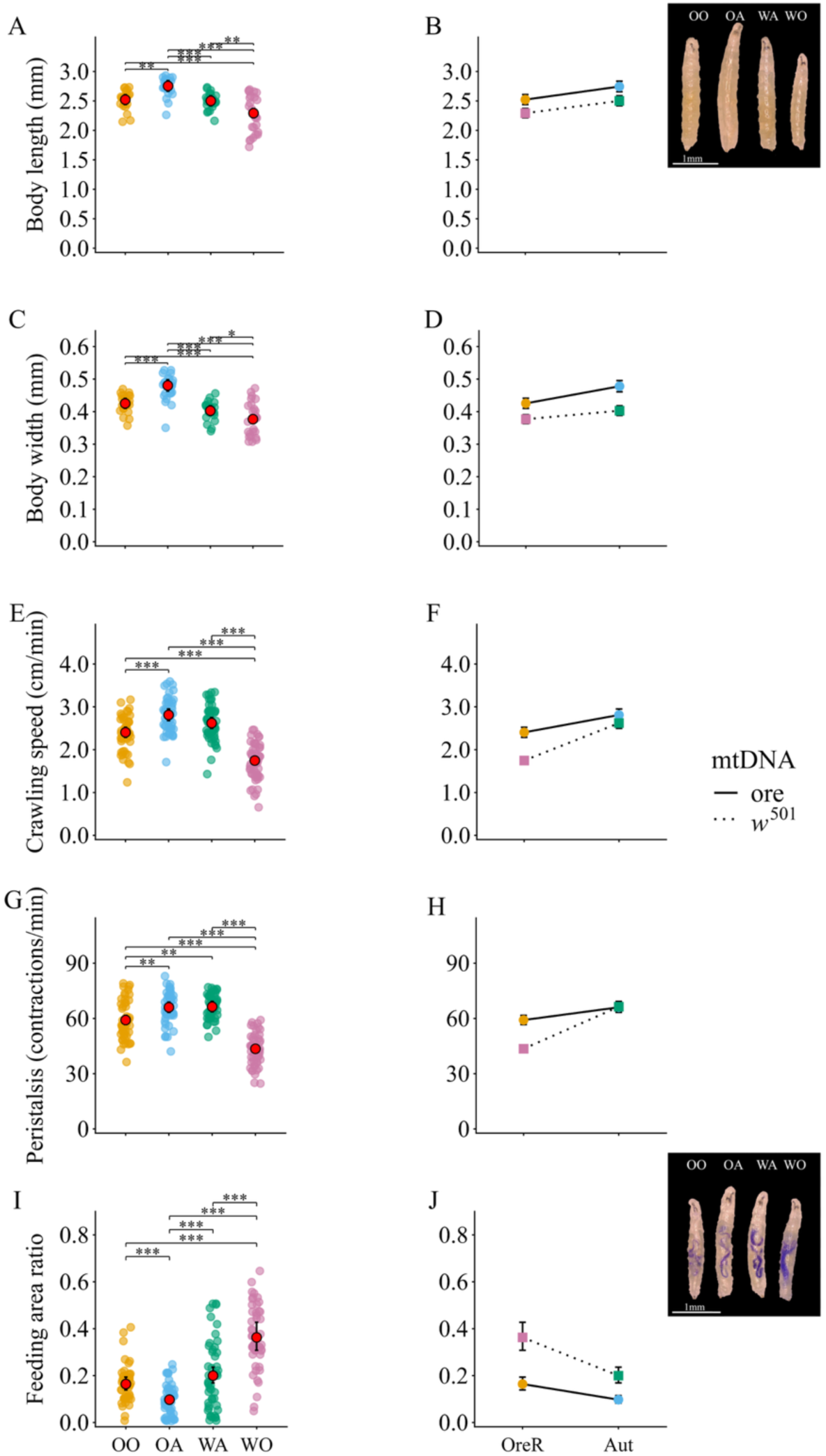
Effects of nuclear and mitochondrial genomes on organismal traits. Larvae with mitonuclear incompatibility ((*w^501^); OreR*) show significantly reduced body size (A-D) and crawling (E-H), with a compensatory increase in feeding (I, J). Estimated marginal means ± confidence intervals are shown as red points; interaction plots using EMMs are shown on the right. Significance: \*\*\**P* < 0.001, \*\**P* < 0.01, \**P* < 0.05.

**Table 1:**
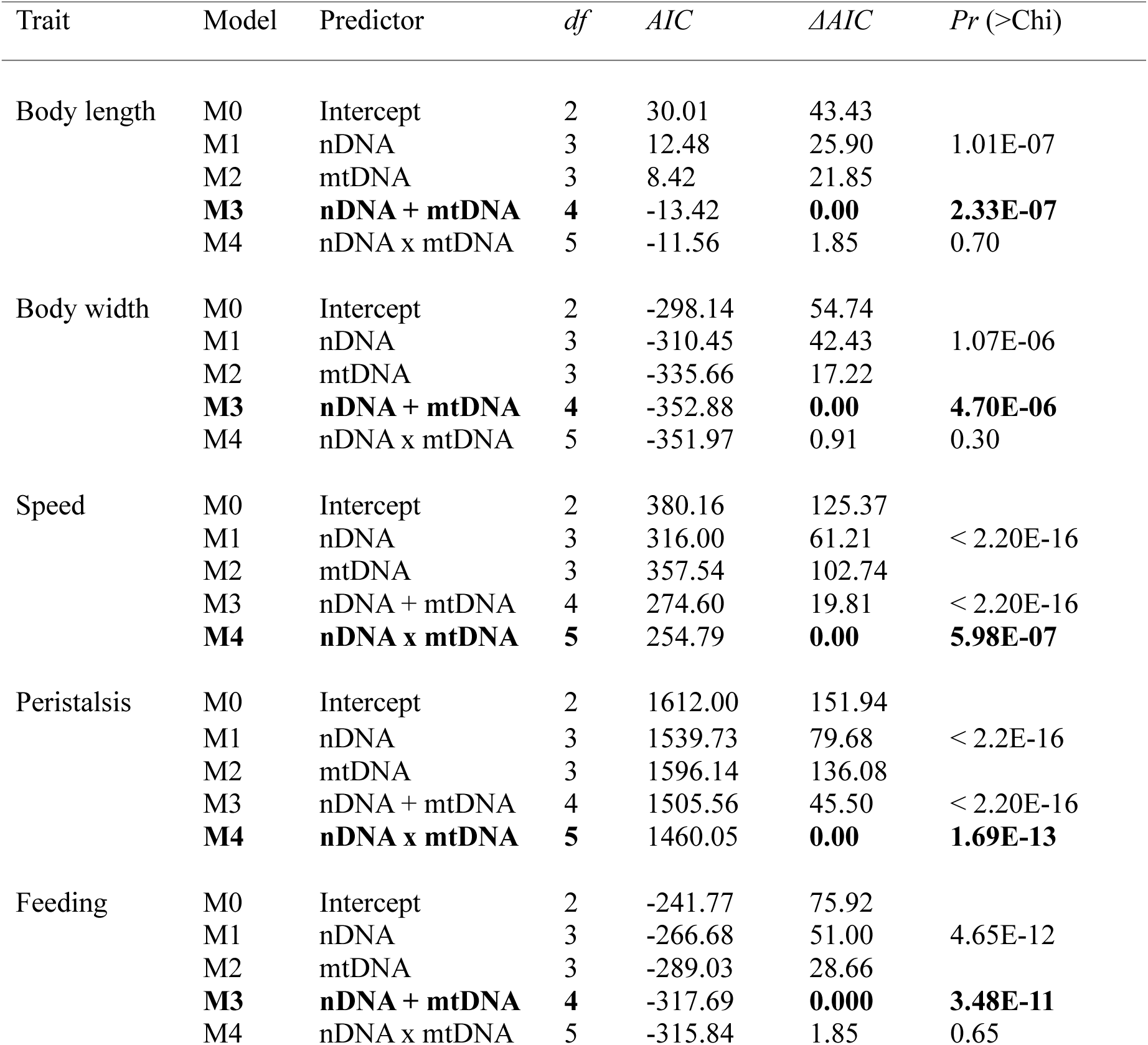
Effect of Nuclear and Mitochondrial Genomes on Organismal Traits. Generalized linear model (GLM) with family gamma and log link were fit for all. ΔAIC values are relative to the best-supported model

| Trait | Model | Predictor | *df* | *AIC* | $\Delta AIC$ | *Pr* (>Chi) |
| --- | --- | --- | --- | --- | --- | --- |
| Body length | M0 | Intercept | 2 | 30.01 | 43.43 |  |
|  | M1 | nDNA | 3 | 12.48 | 25.90 | 1.01E-07 |
|  | M2 | mtDNA | 3 | 8.42 | 21.85 |  |
|  | **M3** | **nDNA + mtDNA** | **4** | -13.42 | **0.00** | **2.33E-07** |
|  | M4 | nDNA x mtDNA | 5 | -11.56 | 1.85 | 0.70 |
| Body width | M0 | Intercept | 2 | -298.14 | 54.74 |  |
|  | M1 | nDNA | 3 | -310.45 | 42.43 | 1.07E-06 |
|  | M2 | mtDNA | 3 | -335.66 | 17.22 |  |
|  | **M3** | **nDNA + mtDNA** | **4** | -352.88 | **0.00** | **4.70E-06** |
|  | M4 | nDNA x mtDNA | 5 | -351.97 | 0.91 | 0.30 |
| Speed | M0 | Intercept | 2 | 380.16 | 125.37 |  |
|  | M1 | nDNA | 3 | 316.00 | 61.21 | < 2.20E-16 |
|  | M2 | mtDNA | 3 | 357.54 | 102.74 |  |
|  | M3 | nDNA + mtDNA | 4 | 274.60 | 19.81 | < 2.20E-16 |
|  | **M4** | **nDNA x mtDNA** | **5** | 254.79 | **0.00** | **5.98E-07** |
| Peristalsis | M0 | Intercept | 2 | 1612.00 | 151.94 |  |
|  | M1 | nDNA | 3 | 1539.73 | 79.68 | < 2.2E-16 |
|  | M2 | mtDNA | 3 | 1596.14 | 136.08 |  |
|  | M3 | nDNA + mtDNA | 4 | 1505.56 | 45.50 | < 2.20E-16 |
|  | **M4** | **nDNA x mtDNA** | **5** | 1460.05 | **0.00** | **1.69E-13** |
| Feeding | M0 | Intercept | 2 | -241.77 | 75.92 |  |
|  | M1 | nDNA | 3 | -266.68 | 51.00 | 4.65E-12 |
|  | M2 | mtDNA | 3 | -289.03 | 28.66 |  |
|  | **M3** | **nDNA + mtDNA** | **4** | -317.69 | **0.000** | **3.48E-11** |
|  | M4 | nDNA x mtDNA | 5 | -315.84 | 1.85 | 0.65 |

#### Crawling

Genotype significantly affected crawling, with *(w^501^); OreR* larvae moving significantly more slowly than the other three genotypes. This pattern was driven by a significant nDNA × mtDNA interaction. GLM-based multiplicative effect sizes indicated that, relative to *(ore); OreR*, *(ore); Aut* larvae were 16.9% faster (exp(β) = 1.16, *P* = 1.15E^-05^), *(w^501^); Aut* were 9.0% faster (exp(β) = 1.09, *P* = 0.01), and *(w^501^); OreR* were ∼27.3% slower (exp(β) = 0.72, *P* < 2E^-16^) (Figure 5E-F, SI T6). Tukey-adjusted pairwise comparisons of EMM showed that *(w^501^); OreR* larvae were significantly slower than both *(ore); Aut* and *(w^501^); Aut* (both *P* < 0.0001). Model comparisons (AIC, ΔAIC, and likelihood ratio tests) supported inclusion of the nDNA × mtDNA interaction (*P* = 5.98E^-07^) (Table 1), with *Aut* nDNA increasing crawling speed relative to *OreR* (*P* < 0.001) across both mtDNA backgrounds (Figure 5F). In contrast, *ore* mtDNA increased crawling speed relative to *w^501^* only when paired with *OreR* (*P* < 0.001), but not with *Aut* nDNA (*P* = 0.18) (Figure 5F). A similar interaction-driven pattern was observed for peristalsis. Relative to *(ore); OreR*, *(ore); Aut* and *(w^501^); Aut* larvae showed 10% (exp(β) = 1.10, *P* = 0.0007) and 11.4% (exp(β) = 1.11, *P* = 0.0004) increases in peristaltic contractions, respectively, whereas *(w^501^); OreR* larvae exhibited ∼26.9% fewer contractions (exp(β) = 0.73, *P* < 2E^-16^) (Figure 5G-H, SI T6). Tukey-adjusted pairwise comparisons of EMM confirmed that *(w^501^); OreR* larvae had significantly fewer peristaltic contractions than both *(ore); Aut* and *(w^501^); Aut* (both *P* < 0.0001). Model comparisons supported inclusion of the nDNA × mtDNA interaction (*P* = 1.69E^-13^) (Table 1), with *Aut* nDNA increasing peristalsis relative to *OreR* (*P* < 0.01) across both mtDNA backgrounds, while *ore* mtDNA increased peristalsis relative to *w^501^* only when paired with *OreR* (*P* < 0.001) and not with *Aut* nDNA (*P* = 0.99) (Figure 5H).

#### Feeding

Genotype significantly affected feeding, and *(w^501^); OreR* larvae fed significantly more relative to the other three genotypes. These differences reflected primarily additive effects of nDNA and mtDNA. GLM-based multiplicative effect sizes showed that, relative to *(ore); OreR*, *(ore); Aut* larvae fed 40.6% less (exp(β) = 0.59, *P* = 1.68E^-05^), *(w^501^); Aut* did not differ (exp(β) = 1.21, *P* = 0.10) and *(w^501^); OreR* fed ∼121.1% more (exp(β) = 2.21, *P* = 2.02E^-10^) (Figure 5I-J, SI T7). Additionally, *(w^501^); OreR* larvae also fed significantly more compared to both *(ore); Aut* (*P* < 0.0001) and *(w^501^); Aut* (*P* < 0.0001) larvae based on Tukey-adjusted pairwise comparisons of EMM. Model comparisons (AIC, ΔAIC, and likelihood ratio tests) indicated that including both nDNA and mtDNA improved model fit (*P* < 3.48E^-11^) (Table 1). *OreR* increased feeding relative to *Aut* nDNA (*P* ≤ 0.001) across both mtDNA backgrounds (Figure 5J). Similarly, *w^501^* increased feeding relative to *ore* mtDNA (*P* ≤ 0.001) across both nDNA backgrounds (Figure 5J). However, the nDNA × mtDNA interaction was not significant (*P* = 0.65).

## Discussion

Our study presents a comprehensive metabolic characterization of mitonuclear interactions in *Drosophila*, providing a systems-level perspective on the contributions from two genomes. nDNA alone had limited effect on metabolome divergence in larvae when expressed with a shared coevolved mtDNA. However, these effects became more pronounced in the presence of a shared but mismatched mtDNA, indicating that mitochondrial background can strongly modulate metabolic phenotype. Larvae with the mismatched *(w^501^); Aut* genotype had metabolome similar to their coevolved control. In contrast, when mismatched genomes exhibited incompatibility in larvae with *(w^501^); OreR* genotype, a more extensive and high-magnitude divergence of the metabolome was observed. In addition, the PCA, hierarchical clustering and RF models consistently separated *(w^501^); OreR* but not *(w^501^); Aut* from the coevolved controls. These findings are consistent with other studies emphasizing that (i) phenotypic outcomes cannot be explained by simple mitonuclear matching or mismatching, and (ii) the effects of mtDNA strongly depend on the nuclear background (Montooth et al. 2010; Joseph et al. 2013; Shastry et al. 2025).

While independent nDNA and mtDNA effects exhibit clear functional modularity in metabolite sets, their epistasis (nDNA × mtDNA) exerted a pleiotropic module-free effect. In general, nDNA was most strongly associated with metabolites involved in core carbon (glycolysis, pentose phosphate pathway (PPP), and carbon storage) and amino acid metabolism (nitrogen and amino acid turnover), and mtDNA with metabolites involved in redox and lipid metabolism (including membrane lipid backbone and fatty acid flux). In contrast, mitonuclear epistasis exhibited a more systemic signature, encompassing metabolites that act both within individual pathways and as intermediates/nodes connecting pathways across metabolic networks. These include metabolites linked to redox-1C coupling, methylation-related processes, lipid remodeling, β-oxidation, nucleotide metabolism, energy currency/redox cofactor pools, central carbon metabolism, and signaling and/or immunomodulatory peptides. Thus, mitonuclear variation captures pervasive non-additive G×G interactions, providing a system to test the omnigenic model (Joseph et al. 2013; Boyle et al. 2017; Dobson et al. 2023), which proposes that trait variation arises from interactions between a few core genes and numerous minute-effect variants across the nuclear genome.

Systemic metabolic remodeling in *(w^501^); OreR* larvae reflects a distinct metabolic trait space primarily geared towards redox and bioenergetic homeostasis, with maintenance metabolism prioritized over larval growth. *Drosophila* larvae increase their body mass over ∼200-fold within 3 days at 25 °C by converting dietary nutrients into biosynthetic intermediates for growth and proliferation (Tennessen and Thummel 2011; Jacobs et al. 2020). In our study (Figure 2 and Figure 6), increased phosphorylated sugars, lactate, polyols and UDP-sugars suggest changes in carbon metabolism, and increased glycolysis for maintenance (Warburg effect). This is consistent with the genetic variation in aerobic glycolysis during L2 that we previously reported in this panel (Matoo et al. 2019). Transcription factors Sima/HIF1α and the *Drosophila* estrogen-related receptor (dERR) have been shown to cooperatively regulate the larval glycolytic program in *Drosophila* (Tennessen et al. 2011; Heidarian et al. 2025). L-2-hydroxyglutarate (L-2HG) also accumulates during normal larval development as part of this metabolic program through a feed-forward mechanism, involving α-ketoglutarate (α-KG), in which lactate dehydrogenase (LDH) promotes L-2HG synthesis, while lactate inhibits mitochondrial L-2-hydroxyglutarate dehydrogenase (L2HGDH)-mediated L-2HG degradation (Tennessen et al. 2011; Li et al. 2017, 2018; Mahmoudzadeh et al. 2020, 2024). L-2HG contributes to hypoxia pre-adaptation and epigenetic regulation, although excessive accumulation is associated with oxidative stress and mitochondrial dysfunction (Li et al. 2017; Mahmoudzadeh et al. 2024). Notably, we found significantly lower L-2HG and α-KG levels in *(w^501^); OreR* larvae, suggesting a distinct biochemical signature under mitonuclear incompatibility that may reflect the altered production and/or consumption of these metabolites. We also observed a broader signature of disrupted nucleic acid homeostasis (accumulation of purines and low pyrimidines), reduced key TCA cycle intermediates (α-KG, fumarate, and isocitrate), lipid remodeling (accumulation of long-chain fatty acids and carnitines, altered fatty acid desaturation– elongation balance and increased phosphatidylcholine and phosphatidylethanolamine), severe amino acid depletion (both glucogenic and ketogenic amino acids, including the essential amino acids) and changes in one-carbon metabolism and/or methylation (elevated S-adenosylmethionine (SAM), with a compensatory activation of antioxidant pathways (elevated glutathione metabolism and other antioxidant metabolites). We acknowledge that our approach provides only a global snapshot of metabolites, and that flux control analyses, targeted metabolomics and characterization of pathway enzyme activities are required to develop a robust mechanistic understanding of this metabolic phenotype.

**Figure 6.**
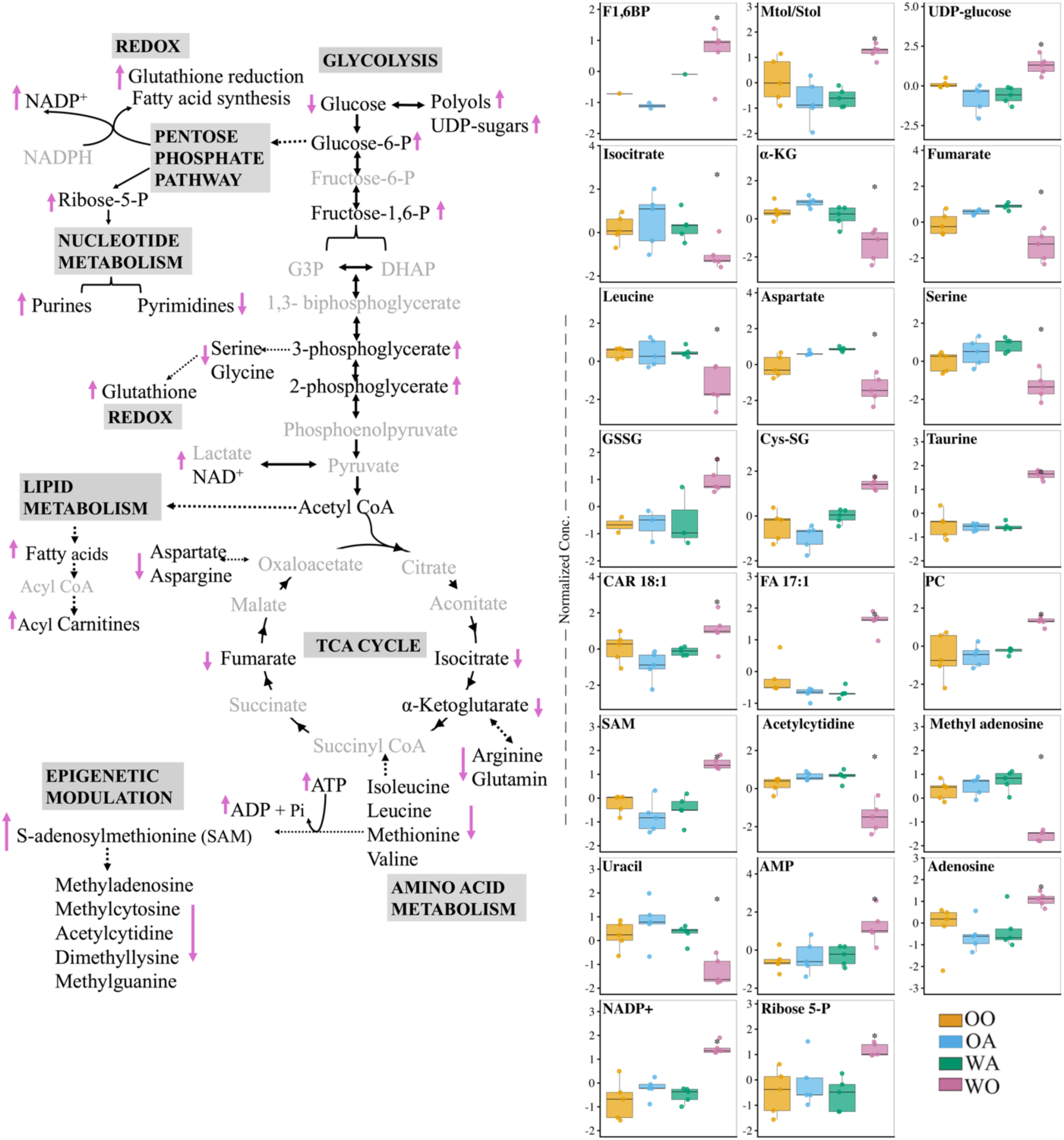
Metabolic rewiring in larvae with mitonuclear genome incompatibility. Larvae with mitonuclear incompatibility ((*w^501^); OreR*) show significantly altered metabolome. (Left) schematic of central carbon, amino acid, lipid, nucleic acid, redox and epigenetic biochemical pathways. Pink arrows represent differentially abundant metabolites in *((w^501^); OreR)* compared to three other genotypes. (Right) Box plots of representative metabolites (*P*-adj < 0.05). F1,6BP: Fructose 1,6 bisphosphate; Mtol/Stol: Mannitol/Sorbitol; GSSG: Oxidized glutathione; Cys-SG: Cysteine glutathione disulfide; CAR18:1: Oleoyl-L-carnitine; FA17:1: Heptadecenoic acid; PC: Phosphorylcholine; SAM: S-adenosylmethionine; AMP: Adenosine monophosphate; NADP+: Nicotinamide adenine dinucleotide phosphate.

Together with delayed growth (Meiklejohn et al. 2013), reduced body size in *(w^501^); OreR* larvae may reflect systemic metabolic rewiring away from biosynthesis. Similar phenotypes occur in the OXPHOS-deficient *Drosophila tko^25t^* strain, which exhibits developmental delay and reduced larval body size (Jacobs et al. 2020, 2023). While the depleted NADPH via pyruvate-malate conversion has been reported in *tko^25t^* (Jacobs et al. 2020, 2023), we speculate that depleted pyrimidines and amino acids, altered adenylate currency, limited TCA cycle might contribute to the size reduction in *(w^501^); OreR* larvae. These changes may affect energy- and nutrient-sensing pathways, including AMP-activated protein kinase (AMPK) and *Drosophila* Target of Rapamycin (*d*TOR), with *d*TOR being constitutively suppressed during *(w^501^); OreR* development (Buchanan et al. 2018). Additionally, fat body has shown to couple mitochondrial bioenergetics, nutrient sensing and larval growth in *Drosophila*. For example, reduced amino acids in fat body suppressed systemic *d*TOR and low nutrition decreased the expression of *Drosophila* insulin-like peptides (dLIP2 and dLIP5) in the fat body, restricting larval growth (Colombani et al. 2003; Agrawal et al. 2016). Fat-body mitochondrial morphology and bioenergetic activity also change during larval development and in response to low nutrient availability, suggesting that mitochondrial metabolism within this tissue may itself contribute to systemic growth regulation (Sriskanthadevan-Pirahas et al. 2022). Manipulating fat-body metabolism through Mitochondrial Transcription Factor A (TFAM) RNAi knockdown or LDH overexpression suppressed the fat-body derived negative regulators ImpL2 and TNF-α/Eiger, thereby enhancing systemic insulin signaling and accelerating larval growth (Sriskanthadevan-Pirahas et al. 2022). Furthermore, whole-body heterologous expression of alternative oxidase (AOX) in *w^1118^ Drosophila* larvae reduced mitochondrial coupling efficiency but increased central carbon flux, thereby accelerating development and suggesting that reduced OXPHOS efficiency can, under some conditions, stimulate rather than limit growth (Garcia et al. 2025). However, AOX in the *tko^25t^* strain, which harbors defects in mitochondrial translation, failed to rescue developmental delay (Kemppainen et al. 2014), indicating that genetic architecture and severity of mitochondrial defect influence growth and developmental phenotypes in *Drosophila* larvae.

Reduced crawling speed and peristalsis in *(w^501^); OreR* are emergent properties of motor neuron activity, muscle contraction, and metabolic state, all of which are sensitive to levels of metabolites involved in mitochondrial bioenergetics (e.g. adenylate pool and fatty acids) and synaptic activity (e.g. amino acids such as glutamine, arginine and aspartate (Stanley et al. 2026)). Previous studies similarly report impaired locomotion in *(w^501^); OreR*, including reduced climbing and pupation height (Meiklejohn et al. 2013) and impaired flight associated with altered mitochondrial morphology in adults (Holmbeck et al. 2015). Finally, *(w^501^); OreR* larvae exhibited increased yeast consumption following starvation, suggesting compensatory nutrient acquisition. Because *Drosophila* naturally rely heavily on yeast as a source of protein and amino acids (Colombani et al. 2003; Buchanan et al. 2018; Jacobs et al. 2020), we speculate that reduced glucose and amino acid availability, mitochondrial dysfunction, and constitutively suppressed *d*TOR could generate an internal starvation-like state that promotes feeding in *(w^501^); OreR* larvae. This interpretation is consistent with previous diet-dependent phenotypes reported in *(w^501^); OreR* and other *Drosophila* models (Mossman et al. 2016; Montooth et al. 2019; Camus et al. 2020; Rodriguez et al. 2021; Bettinazzi et al. 2025). However, the reduced body size suggests that this increased nutrient intake does not translate proportionally into biosynthetic output in *(w^501^); OreR*. In addition to canonical AMPK–*d*TOR, parallel modulation of IIS, Upd2–JAK/STAT, and enteroendocrine gut–brain signaling pathways may also contribute to altered feeding and nutrient demand (Jacobs et al. 2020, Rai et al. 2025). Elucidating the contribution of these candidate pathways under diet-mitonuclear interactions warrants further investigation.

Although our genetic panel is not a natural system, this study emphasizes that genotype-dependent changes in metabolism could have significant evolutionary implications, particularly in populations experiencing genetic introgression, hybridization or rapid and novel environmental shifts. Accordingly, quantifying *post facto* organizational features of metabolism - flux coupling, functional redundancy, modularity, and exaptation (Sambamoorthy et al. 2019) - can help reveal how these evolutionary processes restructure metabolic networks. Whether metabolic rewiring is truly adaptive remains uncertain, but data from our lab and others suggest that mitonuclear mismatch is primarily deleterious. For example, reduced metabolic efficiency, locomotor activity, and body size in *Drosophila* larvae could compromise foraging, predator avoidance, and fitness. Mitonuclear effects have shown to also persist into adulthood, with tradeoffs in immune function, fecundity, and lifespan that are exacerbated by environmental stress (G×G×E) (Hoekstra et al. 2013; Villa-Cuesta et al. 2014; Wolff et al. 2014; Hoekstra et al. 2018; Buchanan et al. 2018; Rand et al. 2022). Because the metabolome is an integrated readout of its genetic, transcriptomic and proteomic variation (Johnson et al. 2016), it serves as a proximal link between the organism and its environment (Bundy et al. 2009). Future studies integrating steady-state metabolomics, flux analyses, and fitness metrics across diverse environmental contexts will be essential to elucidate how metabolic adjustments arising from mitonuclear incompatibility shape metabolic plasticity and influence evolutionary trajectories.

## Data availability

Mitonuclear panel genotypes are available upon request. All supplementary files are deposited in FigShare and will be open access upon publication (private link: https://figshare.com/s/85371816106a2563fc55)

## Acknowledgements

The authors gratefully acknowledge Dr. Kristi L. Montooth for sharing mitonuclear panel genotypes with OBM. We also like to thank her and Dr. Colin Meiklejohn (University of Nebraska-Lincoln) for discussion of data. We acknowledge the use of AI to assist with R script debugging.

